# Brain Organoids, Lessons from Fetal Neocortex Formation, and Rational Design for Quality Control

**DOI:** 10.64898/2026.05.27.728332

**Authors:** Michael Q. Fitzgerald, Teng Zhou, Honieh Hemati, Daniel Labelle, Aiden Momtaz, Erica Harris, Shubham Patil, Gautham Prabhakar, Haonan Hu, Nidhi Pareddy, Visvesh Jegadheesh, Jie Cui, Alysson R. Muotri, Reem Khojah, Shankar Subramaniam

**Affiliations:** Department of Bioengineering at the University of California, San Diego. La Jolla, CA 92093; Department of Pediatrics at the University of California, San Diego. La Jolla, CA 92093

## Abstract

Recent widespread adoption of cerebral organoid protocols has led to many new studies assessing the human-specific features of neural diseases. However, not all organoid studies employ proper quality control, which limits the physiological relevance of their findings. Here, we discuss the stages of in vivo neocortex formation and how those stages are recapitulated in organoid protocols. We then present the first guide for real-time operator removal of maldeveloped organoids in shaking culture. Finally, we show preliminary work on an organoid imaging and mesofluidic control platform for automated quality control of organoid development. Taken together, this approach for assessing the morphological features of organoids will improve the rigor and reproducibility of organoid studies, increase effect sizes of physiologically relevant disease etiology, and pave the way for cortical organoid GMP in high-throughput.

**Clinical Relevance:** This establishes high-throughput, visual brain organoid quality control for disease studies and preclinical testing.

## I. Introduction

The advent of induced pluripotent stem cells (iPSCs) from somatic cells in 2006 [1] ushered in a wave of subsequent methods for the generation of neural stem cells and postmitotic neurons from iPSCs. For the first time, direct perturbation and measurement of patient-specific human neural circuits was possible and widely used for development and disease studies. However, 2D neurons and 3D neurospheres lacked the layered organization of cortical neural circuits observed in vivo (Fig. 1). Then, in 2008, the Sasai Lab, pioneered the first methods for modeling a nascent neural plate with polarized neuroepithelium from stem cells [2]. Over the next decade, more advanced protocols for the maturation of this neuroepithelium into cortical layers emerged in scientific literature. Briefly, all brain organoid protocols utilize supplementation of neural patterning factors that guide the innate differentiation potential of stem cells towards neural fate. Some protocols may utilize many patterning factors and yield cortical-specific organoids, while other protocols rely almost entirely on endogenous genetic programming to generate both cortical and non-cortical regions. If differentiated correctly, all cerebral organoid protocols result in regionally defined neuron layers and a mix of glial support cells surrounding regions of polarized neuroepithelium, similar to the developing neocortex (Fig. 1, bottom). To support 3D growth, some organoid protocols embed individual organoids in ECM-like environments, while shaker-based protocols offer a higher-throughput approach to organoid suspension. Although successfully matured rosettes are comparable between static and shaking cultures, the morphological appearance of neuroepithelia and rosette formation is very different [3]-[9]. Currently, a detailed visual quality control guide for cortical organoids grown in shaker culture systems does not exist.

**Figure 1.**
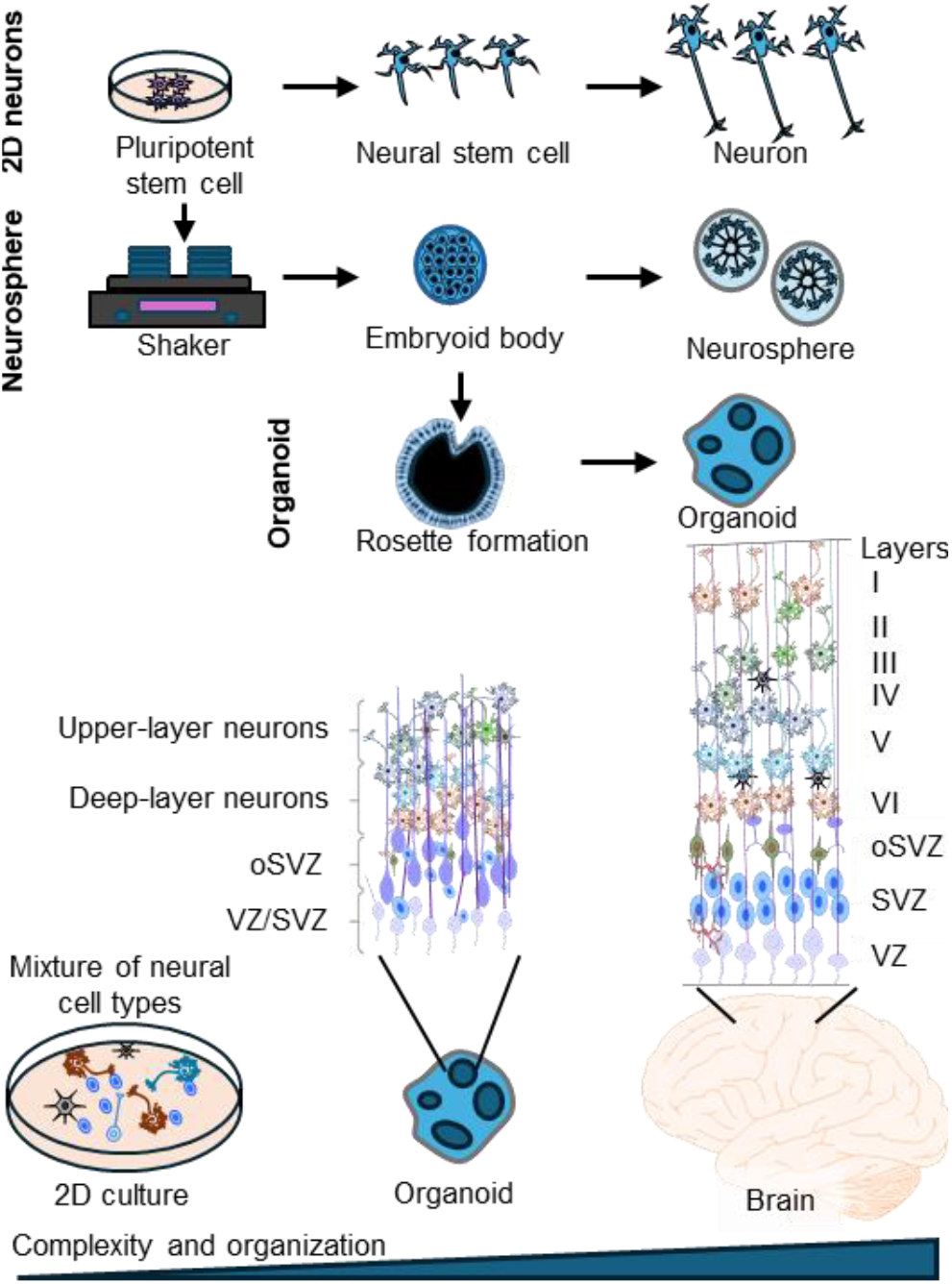
Schematic representation of 2D neuron, neurosphere, and organoid generation (top). The physiological relevance of organoids compared to 2D models and the human brain (bottom). This figure was created using the Motifolio PPT Drawing Toolkits (www.motifolio.com)

Here, we summarize the genetic patterning involved in human cortex development and then overlay this with the factors used in modern organoid protocols to induce cortex formation from stem cells. Understanding the relationship between the differentiation factors added and the expected change in organoid morphology forms the basis for our proposed quality control framework. Therein, proper visual organoid morphology helps verify expected progression of organoid development. Finally, we present preliminary work on dynamic fluidic control and AI-assisted image analysis of organoid development. Application of automated culture and QC to shaker-based culture systems represents a path to high-throughput, GMP-compliant cortical organoid generation.

## II. Brainformation and organoid protocol design

Early morphogen patterning in embryonic blastocysts causes regionally distinct differentiation of stem cells into meso-, endo-, and ectoderm progenitors, which give rise to all somatic cells. Notably, areas of the ectoderm with low FGF, BMP, and TGFβ darken and form the neuroectoderm, which then polarizes and forms the neural tube. The developing neural tube can be divided into three main regions: the forebrain, midbrain, and hindbrain. Most notably, the forebrain gives rise to the cerebrum, which houses cortical (dorsal) and striatal (ventral) structures and is responsible for most thought and movement initiation. The goal of cerebral organoid protocols is to mimic forebrain development with fidelity in vitro. First, pluripotent stem cells are aggregated into 3D spheres called embryoid bodies (EBs). Withdrawal of FGF from EB growth media is sufficient to induce neuralization of embryoid bodies in “unguided” protocols, but further forebrain patterning in “guided” protocols is accomplished via inhibition of TGFβ and BMP, directly mimicking the morphogen conditions that induce forebrain neuroectoderm formation in human gastrula.

In both EBs and the human gastrula, neural plate formation begins when neuroectoderm progenitor cells organize into polarized neuroepithelial cells and expand. In static organoid cultures, neuroepithelial cells align into rosettes locally and bud outward as the neuroepithelia expands [5],[6],[8],[9]. In shaker cultures, the neuroepithelial cells along the entire periphery of the organoid align radially, with an apical-out orientation, and then fold inwards to form ventricles. In either case, the lumen surrounding the new ventricle is called the ventricular zone (VZ) (Fig. 2a, left). In both the neural tube and organoids, the VZ maintains the cell bodies of mitotic neuroepithelial cells at its apical edge and functions as a source of neurogenesis at its basal edge [10]. In dorsal forebrain organoid protocols, the expansion of regionally defined cortical layers is possible with effective maturation of the newly formed ventricular zone (Fig. 2a). For maturation to occur, ventricular radial glia (vRG) switch from symmetric division to asymmetric divisions, which produces one vRG and one neural progenitor cell at the basal edge of the VZ (Fig. 2b, neurogenesis). This region of intermediate progenitors and radial glia is referred to as the subventricular zone (SVZ). This SVZ can go on to generate neurons with cortical plate identity, but production of outer radial glia (oRG) from vRG is required for large and regionally distinct cortical structures. oRG accumulate and form the outer SVZ (oSVZ, Fig. 2b). Both vRG and oRG extend basally to facilitate cortical plateb expansion, but while vRG maintain contact with the apical ventricle, oRG are anchored to the oSVZ and span the entire cortical plate throughout its formation. Beyond this, oRG can also facilitate gliogenesis [11]. To begin cortical plate formation, oRG and intermediate progenitor cells generate immature neurons in the SVZ and oSVZ that then migrate along radial glia and differentiate first into deep-layer, then upper-layer neurons (Fig. 2b, right). In organoids, most protocols utilize a variety of neuronal support supplements to initiate cortical plate formation, including B27, BDNF, NT3, and GDNF. In most protocols, the cortical plate finishes expanding at around month four to five (Fig. 2a, right), which aligns with the in vivo development of the cortical plate between weeks 5 and 20 in vivo [12]. Further maturation of the cortical plate is possible with sustained organoid culture with B27 supplement. Additionally, some protocols switch to basal media with a lower osmolarity to better replicate the osmolarity of the developing neocortex, which decreases with time and in turn makes it easier for action potential depolarization to occur. Interestingly, “semi-guided” cerebral organoids that generate both excitatory and inhibitory neurons display an emergence of complex oscillatory waves between 6 and 10 months that matches the emergence of neural oscillations captured with EEG measurements of preterm neonates [13],[14].

**Figure 2.**
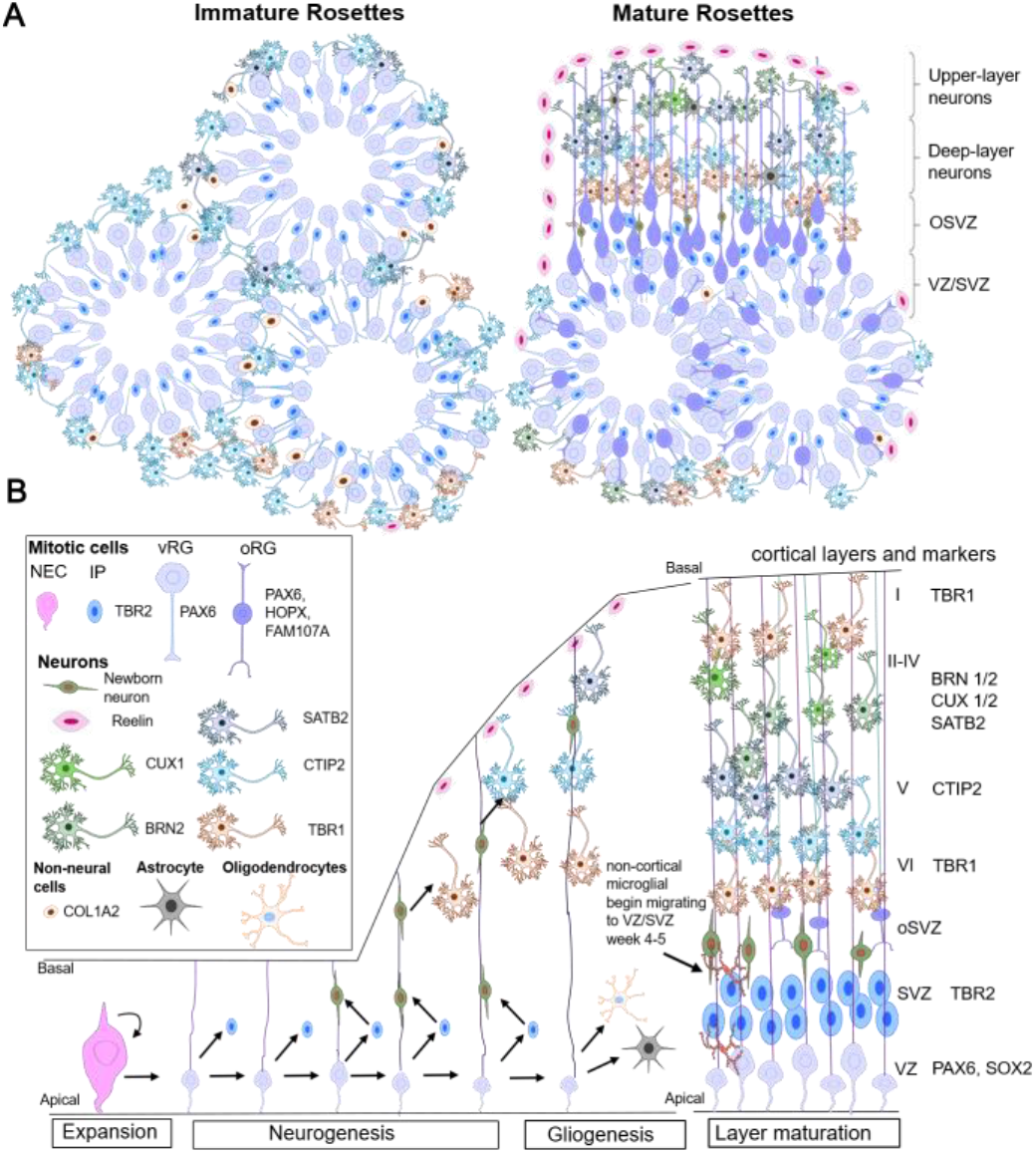
Overview of neuroepithelial infolding and cortical layering. A)Schematic comparison between immature and mature rosettes. B)Developmental progression of cortical layering with marker genes. This figure was created using the Motifolio PPT Drawing Toolkits

## III. Rational design of quality control

Previous methods to model neural circuits fail to recapitulate cortical layering or the presence of human-specific support cells, like oRG (Fig. 1). However, modern three-dimensional organoid protocols generate an expanded neuroepithelium and oSVZ that give rise to defined cortical layers (Fig. 2a, right). Thus, cortical organoids have been widely adopted for disease study. However, these features are dependent on effective neuroepithelium and rosette formation. While organoids must be harvested to confirm the presence of cortical layers, early visual quality control can be performed to ensure successful neural induction, neuroepithelial expansion, and rosette formation. Given that up 1,800 EBs can undergo neural induction on a single 6-well plate in a shaking culture, it is important to pay close attention that EBs properly develop into organoids instead of neurospheres. Here, we present the morphological stages associated with culturing either “guided” or “semi-guided” cortical organoids in a shaker at 95RPM. These protocols were previously developed in the Paşca and Muotri laboratories respectively[13]-[16], with the Paşca protocol being adapted here for shaking culture from day 1.

To generate three-dimensional organoids, iPSCs are first aggregated into embryoid bodies (EBs) using homogeneous microwells. Here, 3 million iPSCs were seeded per well of an AggreWell-800 plate, yielding up to 300 EBs per well that were then transferred to one well of a 6-well plate. Preparation of highly pluripotent and mitotic iPSCs should yield defined aggregates 250-500 μm in diameter with minimal cell death in the AggreWell (Fig. 3, row 1). Inducing forebrain neuroectoderm with FGF withdrawal and TGFB, BMP, and Wnt inhibition darkens the EB and limits mitosis, similar to neuralization in vivo. Therefore, at the end of neural induction, EBs should be dark and 250-500 μm in diameter (Fig. 3, row 2). Following neural induction, FGF and EGF supplementation induce the formation and expansion of cortical neuroepithelial tissue (Fig. 4, row 3). Once the tissue polarizes, the apical edge of the neuroepithelium will be facing outwards and, after expanding, infold to form large VZs with correct apical-basal polarity (Fig. 4, row 4). After about three weeks of development, organoids should be dark, at least 500 μm in diameter, and exhibit obvious neural rosettes. At this stage, the ventricular zone has expanded and begun asymmetric division of vRG to form neural progenitor cells and a defined SVZ. Once organoids are switched to neural maturation media, cortical plate formation begins (Fig. 3, row 5). At the basal end of the SVZ, exogenously supplemented BDNF and NT-3 trigger maturation of progenitor neurons, which in turn migrate along radial glia to form cortical layers. While BDNF and NT3 are only added for a couple of weeks, cortical layering continues over the course of months, and the observable changes by the operator should include 1) a modest increase in organoid size while still retaining dark color (Fig. 3, row 6) and 2) a decrease in media conditioning as neurons switch to oxidative energy metabolism. During this time, about twenty to fifty 1-2mm organoids can be cultured in one well of a six well plate with minimal maintenance. Organoids should be redistributed across wells as they grow as to not limit growth, over-condition media, or aggregate, as previously described [13].

**Figure 3:**
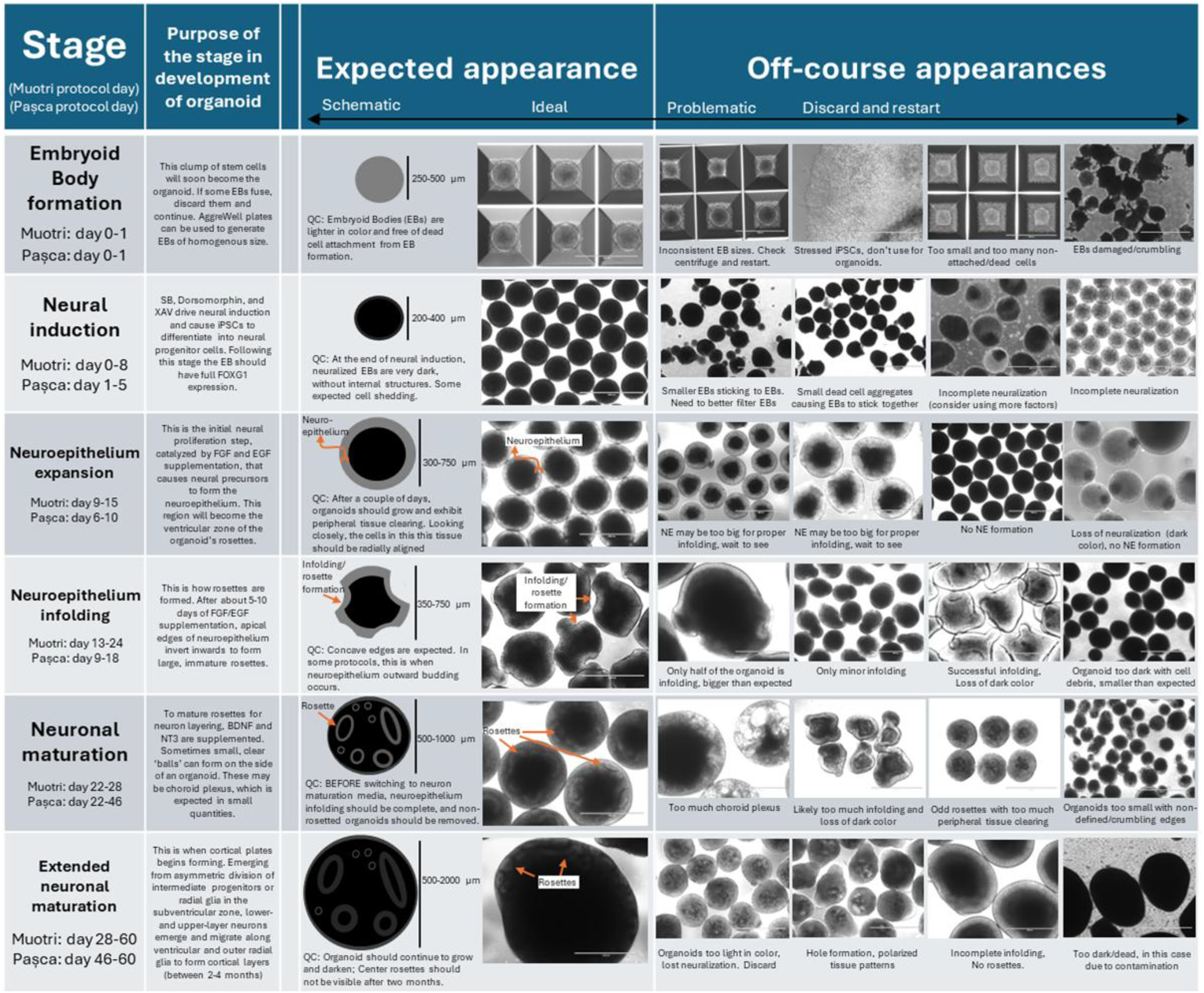
Quality control guide for cortical organoid generation using a shaker culture system. Muotri protocol was performed as described in [13] and Paşca protocol was performed as described in [16], except 1) EBs were cultured in 6-well plates on a shaker at 95rpm from day 1, and 2) LIF was added during neuroepithelial expansion steps. Scale bar: 1000 μm

**Figure 4.**
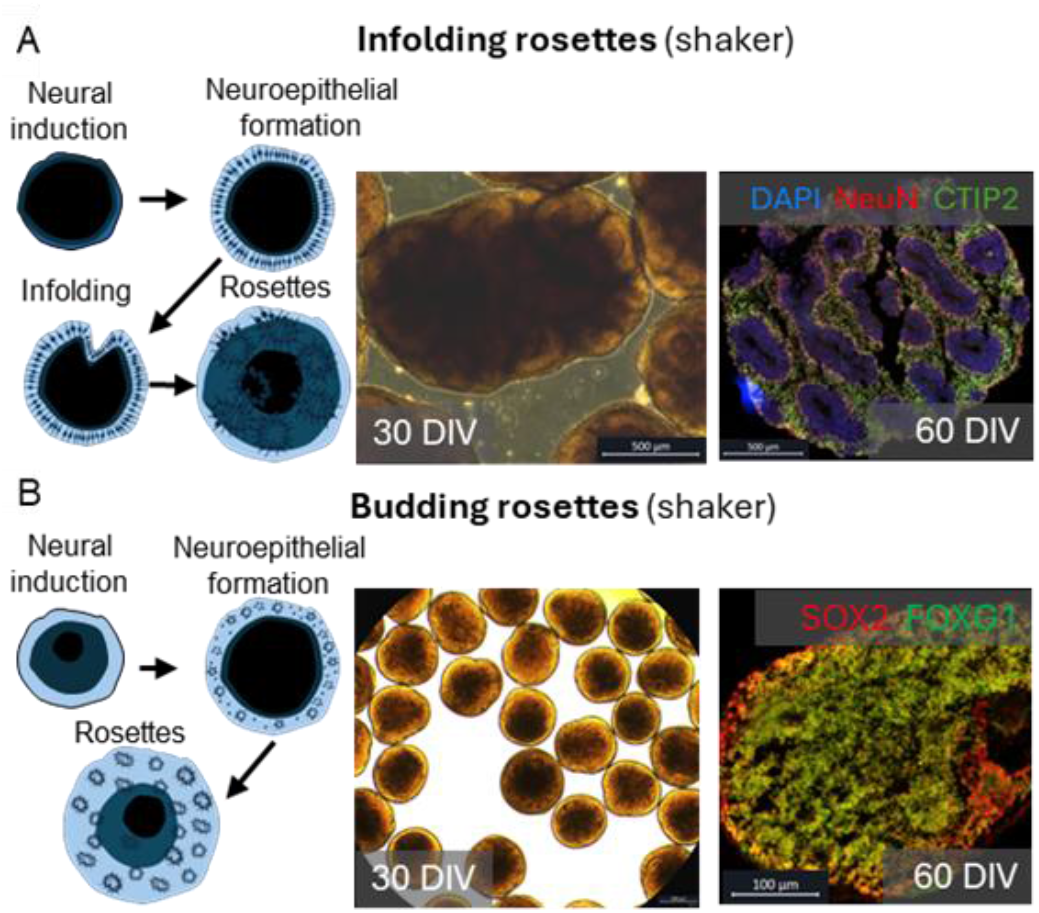
Overview of neuroepithelial infolding and cortical layering. (A) schematic showing the apical-out orientation of organoids grown in shaking culture, with subsequent infolding for rosette formation. Brightfield and immunostaining images show large rosettes, scale bar: 500μm. B) Schematic showing smaller, internally-formed rosettes in shaking culture. Brightfield and immunostaining images show large rosettes. scale bar: 500μm and 100μm, respectively. This figure was created using the Motifolio PPT Drawing Toolkits (www.motifolio.com)

As a standard of practice, immunostaining for oRG, specific separation of upper- and lower-plate markers (SATB2 and TBR1), and forebrain (FOXG1) should be performed in a batch-by-batch manner to ensure presence of these structures, as many have done previously [11],[17]. For single cell sequencing studies, the Human Neural Organoid Cell Atlas [18] is available to allow direct mapping of cortical organoids for cell type and regional labeling (Supp. Fig. 1). The cell type profile generated helps ensure correct cell composition and regional identity of the organoid. Failure to obtain these features should beget changes in subsequent organoid preparation. For example, a new study suggests that LIF can facilitate oRG formation and that organoid researchers should supplement LIF during neuroepithelia formation if oRG or cortical layering is not observed from properly formed rosettes [11]. Alternatively, if organoids do not have FOXG1+ forebrain identity, then the supplied neural induction factors should be reassessed as fore-to hindbrain specification occurs during neuroectoderm development. While all studies should incorporate these end-stage quality controls, real-time quality control of early stages of organoid development is easy and can be done by any operator with a tabletop microscope and Figure 3. Variation associated with generating cortical organoids is anticipated, which is why pruning of non-rosetted or incorrectly rosetted organoids is required.

Of note, while organoids will infold multiple times, the human neural tube only closes once. Current efforts in cortical organoid reproducibility and physiological relevance aim to generate organoids with a single ventricular zone, with mixed success [19]. In lieu of this, neuroepithelial formation in cortical organoid protocols can be modulated with either FGF exposure or changing the organoid’s physical environment. FGF addition to forebrain neuroectoderm not only aids in cortical specification but also in the expansion of neuroepithelial cells and extension of neuroepithelial cells into ventricular radial glia. Alternatively, embedding EBs in ECM during neuroepithelium formation results in internally formed rosettes with correct apical-basal polarity and facilitates outward neuroepithelia budding [8],[9]. Similarly, individual organoids can be grown in 96-well plates or static low-attachment dishes, which also results in neuroepithelial budding, but at a slower rate [3],[7],[16]. Commercially available kits can be purchased with quality control guides for embedding- and low attachment plate-based organoid protocols [5],[7]. However, these guides are often incorrectly applied to shaker-based cultures and do not mention rosette infolding. Of note, incomplete neural induction can result in internal-born rosettes. However, budding in shaker-based cultures results in significantly smaller rosettes compared to infolded rosettes and should be avoided (Fig.4). If neuroepithelial budding is desired, culturing organoids statically during rosette formation is ideal.

## IV. Engineering quality control for high-throughput

While pruning of individual organoids is the current standard of practice, this process can be time-intensive, require additional operator training, and can ruin experiments if done incorrectly or not at all. Ideally, ensuring uniform progression through development and selection of organoids in real time would be automated and unbiased, especially when organoids are used in disease studies and GMP settings. Therefore, we aimed to design an organoid culture system capable of directly manipulating and imaging organoid cultures in 6-well plates.

As organoids grow from micro-to mesoscale sizes, they face stronger shear and inertial wall lift forces, which limit automated flow control using conventional microfluidic channel and device designs. To address this, we coupled imaging with unconfined, motor-driven, mesoscale flow control in a channel-less setup, allowing direct manipulation of organoids in a closed dish design and identification of features linked to organoid development. First, we performed a parametric study using COMSOL Multiphysics on mesoscale flow dynamics (e.g. velocity and shear stress) as a function of Reynolds number in different motor configurations in one well of a six-well plate (Fig. 5a). We found that Re 50 achieves cytocompatible shear stress levels. We then optimized our parameters for motor setup, depth, and velocity with direct testing on organoids. This resulted in effective herding of organoids to the center of the dish for imaging (Fig. 5b). Additionally, the device is fabricated with 3D printed material (PETG) and humidity and heat-resistant stepper motors and sensors, making it compatible to run inside an incubator for long term (Fig. 5C). Compared to a traditional microscope that moves the stages, our device moves the camera in the XY-plane while keeping the culturing plate at a steady position. A two-week trial was performed to validate the motors and the camera, the deviation from the center of the plate was calculated to be ±150 um (n=29). For future improvement, we aim to implement an objective turret, allowing the device to image organoids at higher magnification. AI-based segmentation YOLOv5 and YOLOv8 was applied to track the size of one-month-old organoids over the course of a week (Fig. 5d-f) and identify wells with organoids that died during this period (Fig. 5c, wells 4 and 6). Validation against manual scoring (Fig. 5e) showed YOLOv8 aligned more closely with manual counts than YOLOv5, with fewer false negatives, though both models undercounted organoids at image edges. Next, we aim to apply this method to organoids during the neuroepithelium expansion phase to identify organoids that exhibit expected growth during this stage. An exciting next step includes a more complex AI-based analysis of organoid morphology and cross-referencing with Fig. 3 to ensure rosette formation and give organoids a quality control score at each stage of development. Finally, this could be coupled with refined motor systems for removal of low-quality organoids and, thus, enable end-to-end autonomous quality control of organoid development.

**Figure 5.**
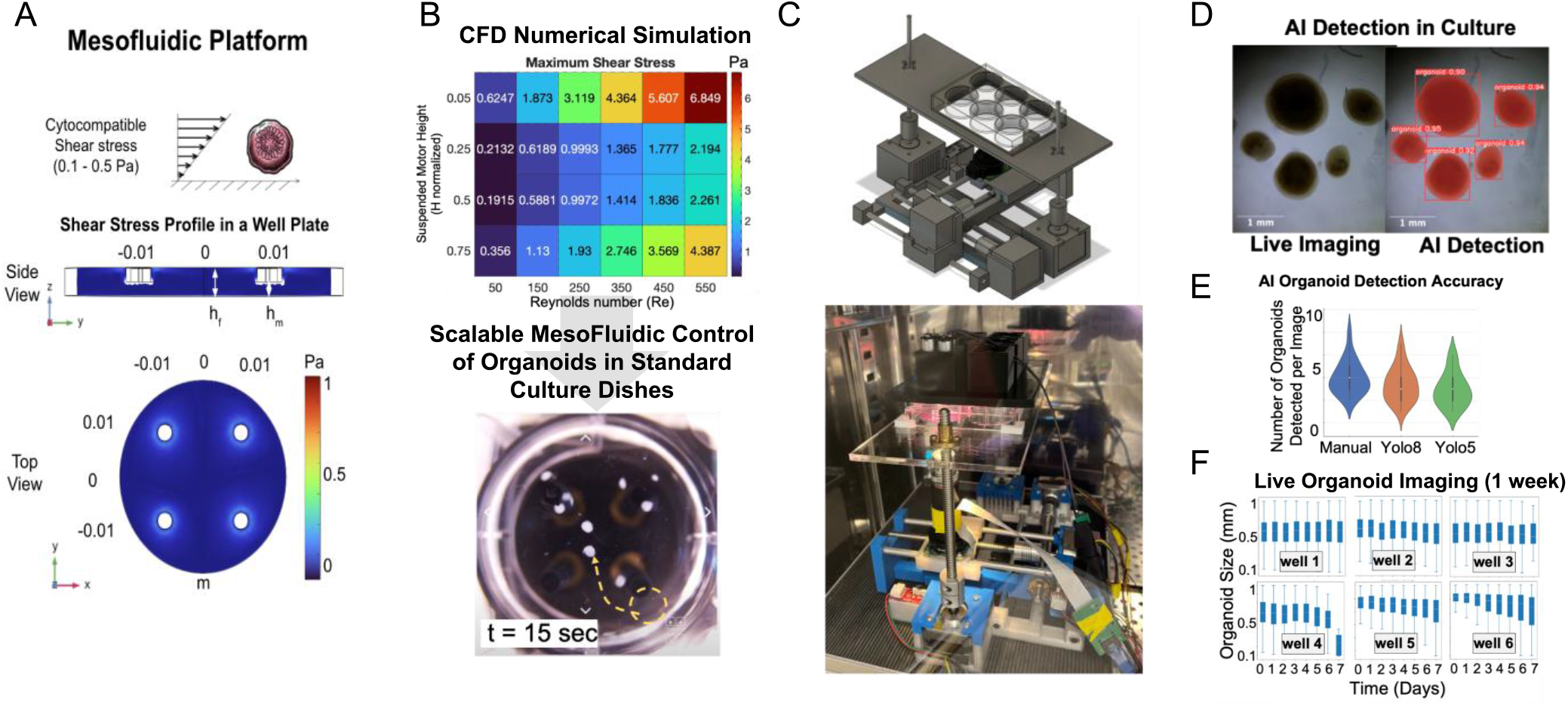
Schematic for automatic imaging and analysis system. A) Top and side view of the velocity profile of the CFD simulation. Motor gaps are 13mm, motors are submerged .75mm, and Re equals 550. B) CFD optimized setup based on motor setup and Reynold’s number and demonstration of moving cultured organoids C) show the CAD design of the camera system and the experimental setup running inside the incubator. D) Image of organoids in culture and overlay of AI-based image segmentation. E) Violin plots comparing manual annotation and the two AI detection models. F) Box and whisker plots for organoid sizes in each well over the course of a week

## V. Conclusion

Seemingly innocuous, a simple sheet of neuroepithelial cells carries the capacity to generate the body’s most complex tissue–the cerebral cortex. With modern protocols, only a few key steps are needed to generate neuroepithelium and mature that epithelium for cortical plate generation. Unfortunately, almost every step of this process can be influenced by disease. For example, issues with establishing neuroepithelial polarity are linked to schizophrenia, autism spectrum disorder (ASD), and down syndrome [20]. Thus, a clear description of how these structures look during development is warranted, and organoids should be checked for presence of these structures *before* experimentation. In turn, each step of organoid differentiation can also be affected by incorrect lab practices or unexpectedly spoiled reagents. Without informed quality control, organoids with maldeveloped rosettes, or no rosettes at all, can confound entire disease studies. However, real-time organoid pruning represents an opportunity to improve experimental rigor and reproducibility, increase resolution of disease phenotypes, and help bridge the gap between preclinical studies and translatability to human patients. We present Fig. 3 to facilitate this process in shaker-based cultures, which are significantly higher throughput and involve less direct organoid manipulation than other methods. Additionally, it should be noted that organoids with proper rosette infolding still might not develop mature layers. As such, immunostaining and electrophysiology should be performed to confirm the presence of active cortical layers. Further, observed morphology will change based on the protocol used. Timing and concentration of factors should be adapted to ensure rosette infolding.

Finally, we present a proof-of-concept system for autonomous organoid imaging and analysis that can be used for certain aspects of quality control. In the future, we aim to expand the capabilities of this system for end-to-end autonomous organoid quality control based on Fig. 3. In the meantime, we encourage the adoption of rosette-based quality control coupled with microscope-assisted organoid pruning by the operator for real-time quality control.

## Author contributions

M.Q.F. wrote the manuscript, Fig. 1, 3, and 4, and contributed to research design for Fig. 5. M.Q.F. and H.H. differentiated organoids and took images for Fig. 3. D.L. assisted organoid culture, expanded the rough draft manuscript and added commentary on human specific features of cortical organoids. E.H. and M.Q.F. generated Fig. 2. T.Z., S.P., H.H, G.P, N.P, V. J., J.C. Performed experiments for fig. 5. T.Z. and M.Q.F. generated organoids for Fig. 5. R.K. oversaw research design and experiments for Fig. 5. A.M. performed alignment annotation for supplemental Fig. 1. S.S. and A.R.M. were involved in overall research design and supervision.

## Acknowledgment

Thank you to Angels Almenar and Miguel Tenreiro for mentorship and support.

## Ethics Approval

The experiments using iPSCs in this study were approved by the University of California San Diego Institutional Review Boards and Institutional Stem Cell Research Oversight Committees guidelines and regulations (no. 141223ZF)

## Supplementary figure

**Supplemental figure 1.**
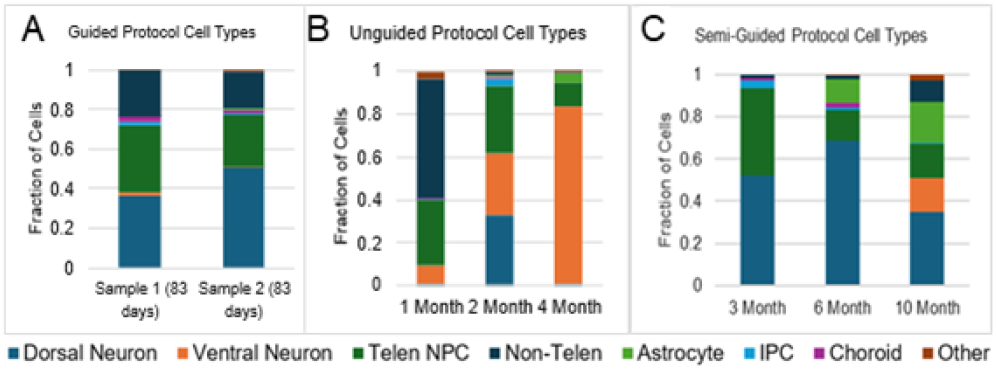
Cell type composition across organoid protocols using Human Neural Organoid Cell Atlas [18]. A) Guided cortical protocol from the Paşca lab [21] exhibit almost entirely dorsal identity. B) Unguided cerebral organoids generated from exhibit dorsal and ventral telencephalon identities [22]. C) Semi-guided cortical organoids generated from the Muotri lab [13] exhibit both dorsal and ventral identities. scRNA-seq datasets were preprocessed using standard workflows, including total count normalization to one million counts per cell, log-transformation, and filtering out cells with fewer than 200 expressed genes, genes expressed in fewer than 3 cells, and cells with greater than 0.08 mitochondrial gene content to remove low-quality cells and potential debris. Cell type annotations were transferred from the Human Neural Organoid Cell Atlas (HNOCA) by training an SCVI/SCANVI model on the reference data and applying weighted k-nearest neighbor mapping to align and classify query datasets.

## Notes

* Research supported by NIH RF1-AG084030 (SS), Joan and Irwin Jacobs Endowment (SS), and NSF grant No. 2321122 (RK)

### Competing Interest Statement

M.Q.F., S.S., and A.R.M are inventors on a provisional patent involving QC and generation of organoids Atty: P14293US00. A.R.M. is a cofounder of and has an equity interest in TISMOO, a company dedicated to genetic analysis and brain organoid modeling focusing on therapeutic applications customized for autism spectrum disorder and other neurological disorders with genetic origins. The terms of this arrangement have been reviewed and approved by the University of California San Diego in accordance with its conflict-of-interest policies. A.R.M. is an inventor of several patents related to human functional brain organogenesis.

